# A conserved *in vivo* burn wound infection model for diverse pathogenic fungi

**DOI:** 10.1101/2024.11.12.623264

**Authors:** Nayanna M. Mercado Soto, Adam Horn, Nancy P. Keller, Anna Huttenlocher, Andrew S. Wagner

## Abstract

Secondary fungal infections represent a major complication following thermal injuries. However, the mechanisms of fungal colonization of burn tissue and how the host subsequently responds to fungi within this niche remain unclear. We have previously reported a zebrafish model of thermal injury that recapitulates many of the features of human burn wounds. Here, we characterize host-fungal interaction dynamics within the burn wound niche using two of the most common fungal pathogens isolated from burn injuries, *Aspergillus fumigatus* and *Candida albicans*. Both *A. fumigatus* and *C. albicans* colonize burned tissue in zebrafish larvae and induce a largely conserved innate immune response following colonization. Using drug-induced cell depletion strategies and transgenic zebrafish lines with impaired innate immune function, we found that macrophages control fungal burden while neutrophils primarily control invasive hyphal growth at the early stages of infection. However, we also found that loss of either immune cell can be compensated by the other at the later stages of infection, and that fish with both macrophage and neutrophil deficiencies show more invasive hyphal growth that is sustained throughout the infection process, suggesting redundancy in their antifungal activities. Finally, we demonstrate that *C. albicans* strains with increased β(1,3)-glucan exposure are cleared faster from the burn wound, demonstrating a need for shielding this immunogenic cell wall epitope for successful fungal colonization of burn tissue. Together, our findings support the use of zebrafish larvae as a model to study host-fungal interaction dynamics within burn wounds.

**Importance:** Secondary fungal infections within burn wound injuries are a significant problem that delay wound healing and increase the risk of patient mortality. Currently, little is known about how fungi colonize and infect burn tissue or how the host responds to pathogen presence. In this report, we expand upon an existing thermal injury model using zebrafish larvae to begin to elucidate both the host immune response to fungal burn colonization and fungal mechanisms for persistence within burn tissue. We found that both *Aspergillus fumigatus* and *Candida albicans*, common fungal burn wound isolates, successfully colonize burn tissue and are effectively cleared in immunocompetent zebrafish by both macrophages and neutrophils. We also find that *C. albicans* mutants harboring mutations that impact their ability to evade host immune system recognition are cleared more readily from burn tissue. Collectively, our work highlights the efficacy of using zebrafish to study host-fungal interaction dynamics within burn wounds.

## Introduction

The World Health Organization (WHO) estimates that around 11 million burn injuries per year occur worldwide, with almost 200,000 resulting in patient death (1). During burn injury, the skin barrier and microbiome are disrupted, and subsequent antimicrobial treatment and immune system dysregulation make these wounds susceptible to microbial infections (2). Burn wound infections (BWIs) are the leading cause of mortality among burn wound patients and add an additional layer of complexity that can delay the healing process and further alter the immune response (3–5). BWIs are primarily caused by bacterial species like *Pseudomonas aeruginosa* and *Staphylococcus aureus* (3, 6). However, fungal pathogens are also a significant cause of infection at the burn wound interface. Indeed, global reports have shown that up to 44% of severe BWIs develop secondary infections from fungal species (5). Once colonized, the risk for the development of systemic infections increases substantially, with mortality rates reported to be as high as 76% once dissemination occurs (7–9). Despite this high prevalence, little is known about how fungi successfully colonize the burn wound tissue to cause disease, nor how the host subsequently responds to invading fungi at this niche. Indeed, most of the available data is based on clinical studies, highlighting the need for a robust infection model to further elucidate these dynamics.

Although fungal BWIs are understudied, *in vitro* and *in vivo* models do exist that have provided initial insights into the host response and therapeutic strategies following disease onset. For example, BWIs in human tissue explants have been recently established as an *ex vivo* model to study the host response to *Candida albicans* and have elegantly shown a role for neutrophils in responding to both burn damage and fungi (8). Animal burn models using mice, rats, pigs and galleria also have been utilized to study drug efficacy following fungal burn wound colonization (10–12). However, these models have yet to be leveraged to extensively study the mechanisms of fungal colonization and host immune response, and many do not easily permit *in vivo* imaging of host-fungal interactions to better characterize disease dynamics. Zebrafish larvae, which are naturally transparent, provide an advantage for imaging infection kinetics *in vivo* (13–15). These vertebrates have a largely conserved innate immune response with humans, and zebrafish larvae have been established as a reliable model to study the inflammatory profile and tissue remodeling processes following burn injury (16, 17). In addition, we have shown that the inflammatory profile of burn wounds can be altered by *P. aeruginosa* colonization and have recently expanded upon the zebrafish burn model to show that thermal injuries can be colonized by the fungal pathogen *C. albicans* (18, 19). However, the mechanisms of fungal colonization and host mediated clearance remain unknown.

Here, we characterize a fungal BWI model in zebrafish using two of the most common fungal pathogens isolated from burn wounds, *C. albicans* and the filamentous fungus *Aspergillus fumigatus* (2, 6, 7, 9, 20). We demonstrate that both colonize the burn wound and are successfully cleared in an immunocompetent host. We found that macrophages and neutrophils play synergistic and partially redundant roles in controlling fungal burden and hyphal formation, and that infection in zebrafish with both deficient macrophages and neutrophils results in extensive colonization and invasive hyphal growth from the site of injury. We also show that fungal mutants with increased β(1,3)-glucan exposure are cleared faster from the burn site, demonstrating a need for shielding immunogenic epitopes in the fungal cell wall for persistent colonization to occur. Collectively, our data demonstrate conserved immune responses against two distinct fungal pathogens in the burn wound and highlight the potential of zebrafish larvae as a model to study fungal BWIs.

## Results

### Pathogenic fungi colonize burn wounds in immunocompetent zebrafish larvae

We have recently expanded on a preexisting burn wound injury model using 3 days post-fertilization (dpf) larvae to show that *C. albicans* can successfully colonize burn tissue in this model (19). However, characterization of the infection process and the subsequent host response has yet to be assessed. Furthermore, it is currently unknown if additional pathogenic fungi commonly isolated from BWIs are capable of infecting damaged tissue in a larval zebrafish burn model. To address this, we induced burn wound injury in 3 dpf zebrafish larvae via cauterization and incubated fish with either fluorescent labelled spores of *Aspergillus fumigatus* or *Candida albicans* yeast to permit injury colonization (Fig. 1A). Both confocal microscopy images and plating for colony forming units (CFUs) showed that *A. fumigatus* was able to successfully colonize the burn wound, with peak fungal burden at the 24-hour time point followed by dramatic fungal clearance by 72 hours post burn (hpb) (Fig. 1B-E). Importantly, the infection was restricted to the burn wound area by 24 hpb, highlighting the need for tissue disruption for colonization to occur (Supplemental Fig 1D). In addition to colonization, germinated spores and/or hyphae were seen across all time points but was limited to small hyphal extensions localized to the tail fin (Fig. 1D). It should be noted that at time points prior to 24 hpb there is an abundance of fungi superficially adhered to necrotic tissue of the burn wound, but by 24 hpb fungal cells are interstitial within viable tissue. Therefore, we considered the 24 hpb time point as the start of stable colonization of the burn wound for further analysis.

**Figure 1.**
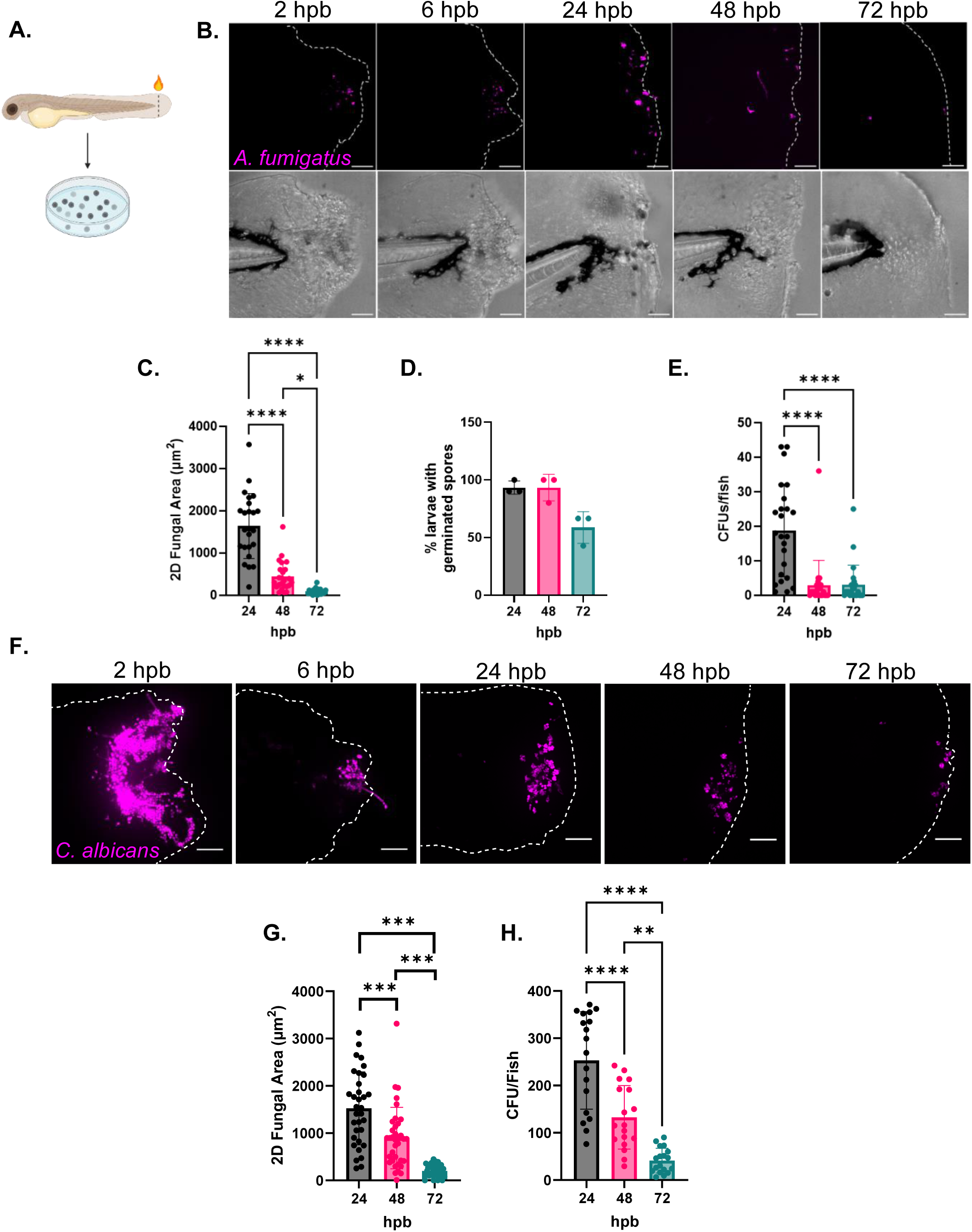
Immunocompetent fish clear fungi from the burn wound. (A) Fungal burn wound infection schematic. The tail fins of 3 days post-fertilization (dpf) larvae are injured using a cauterizer and immediately transferred to E3 -MB media with 2 x 10^7^ fungal cells/ml. (B) Representative images of wild-type larvae infected with fluorescent (RFP) *A. fumigatus* CEA10 and imaged at 2, 6, 24, 48 and 72-hours post-burn (hpb). Images represent maximum intensity projections of z-stacks (scale bar is 50µm). Dashed white lines outline tail fin edge. (C) 2D-fungal (RFP) area from 24-72 hpb. (D) Percent of larvae with germinated spores from 24-72 hpb. Each dot represents percent of an individual experiment. (E) Larvae infected with *A. fumigatus* CEA10 were homogenized and plated for CFUs at 24-72 hpb. (F) Representative images of wild-type larvae infected with fluorescent (RFP) *C. albicans* and imaged at 2, 6, 24, 48 and 72-hpb. Images represent maximum intensity projections of z-stacks (scale bar is 50µm). Dashed white lines outline tail-fin edge. (G) 2D-fungal (RFP) area and (H) CFUs/fish from 24-72 hpb. (I-J) 2D-fungal area at 24 hpb from wild-type larvae innoculated at 3dpf with either live or paraformaldehyde (PFA)--fixed (I) *A. fumigatus* CEA10 spores or *C. albicans* yeast cells (RFP). Bars on graphs presented as mean + standard deviation. Each dot represents data from an individual fish and results represent data pooled from 3 independent experiments. n = 24-30 larvae per condition. *p* values calculated by ANOVA with Tukey’s multiple comparisons. \**p*<0.05, \*\*\**p<*0.001, \*\*\*\**p*<0001.

The ability to colonize the burn wound was conserved by *C. albicans* (Fig. 1F-H). Strikingly, *C. albicans* seemed to persist longer, with higher fungal burden compared to *A. fumigatus,* especially at 72 hpb (Fig 1G-H). Similar to *A. fumigatus*, morphological transition to hyphae was seen during *C. albicans* infection, but the kinetics of these transitions differed. *A. fumigatus* germination was seen starting at 24 hpb, but *C. albicans* hyphae was observed as early as 2-6 hpb and was largely controlled by 24 hpb (Fig. 1F). This is consistent with previously published work in which *C. albicans* germination starts within 6 hours-post infection (hpi) in a hindbrain ventricle zebrafish infection model, but by 24 hpi the host immune system effectively controls hyphal growth (21).

### Fungal colonization increases immune cell recruitment to burns

In both humans and fish, macrophages and neutrophils are recruited to the burn wound and function to clear cellular debris from damaged and necrotic tissue (16, 18). To assess if fungal BWI further impacted immune cell recruitment we burned and infected larvae with fluorescently labeled macrophages (*mpeg-GFP*) and neutrophils (*lyz-BFP*) and quantified leukocyte recruitment. By 24 hpb, macrophages and neutrophils are recruited to the burn wound during both *A. fumigatus* (Fig. 2A-C) and *C. albicans* (Fig. 2D-F) infection, and their numbers drop over a 72-hour period in accordance with wound healing and fungal clearance. Like the inflammatory profile of the burn wound at basal levels (16), macrophages were the predominant cell type present during both fungal infections (Fig. 2B&E). Interestingly, at 24 hpb fish infected with *C. albicans* showed increased levels of both macrophages and neutrophils at the tail fin when compared to uninfected burned fish (Fig. 2E-F), but only neutrophil levels were increased at 24 hpb during *A. fumigatus* infection (Fig. 2C). However, neither *C. albicans* nor *A. fumigatus* significantly impaired tail fin regeneration compared to the uninfected control following burn (Supplemental Fig. 1B-C).

**Figure 2.**
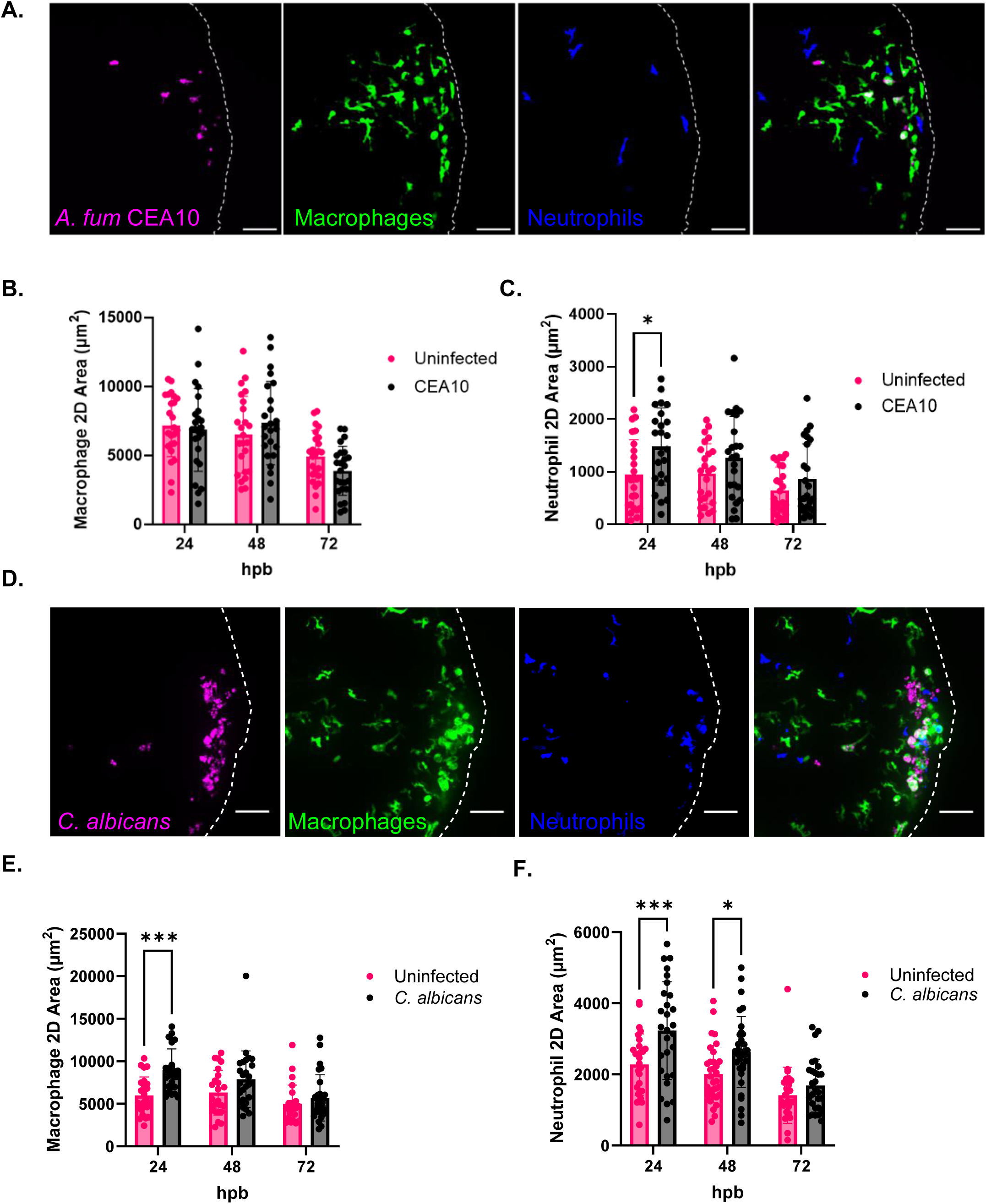
Fungal presence increases immune cell infiltration following thermal injury. (A-C) 3-days post fertilization wild-type larvae with fluorescent macrophages (*mpeg-GFP*) and neutrophils (*lyz-BFP*) were burned and infected with fluorescent *A. fumigatus* CEA10 (RFP). Control larvae were injured but uninfected. Larvae were imaged at 24-, 48- and 72-hpb. (A) Representative images of larvae infected with *A. fumigatus* CEA10 at 24 hpb. Images represent maximum intensity projections of z-stacks (scale bar is 50µm). Dashed white lines outline tail-fin edge. (B) 2D-macrophage area from 24-72 hpb. (C) 2D-neutrophil area from 24-72 hpb. (D-F) 3-days post fertilization wild-type larvae with fluorescent macrophages (*mpeg-GFP*) and neutrophils (*lyz-BFP*) were burned and infected with fluorescent *C. albicans* (RFP). (D) Representative images of larvae infected with *C. albicans* at 24 hpb. Images represent maximum intensity projections of z-stacks (scale bar is 50µm). Dashed white lines outline tail-fin edge. (E) 2D-macrophage area from 24-72 hpb. (F) 2D-neutrophil area from 24-72 hpb. Bars on graphs presented as mean + standard deviation. Each dot represents data from an individual fish and results represent data pooled from 3 independent experiments. n = 24-30 larvae per condition. *p* values calculated by a two-way ANOVA with Šidák’s multiple comparisons test. \**p*<0.05, \*\*\**p<*0.001.

To elucidate which phagocyte may be predominantly driving fungal clearance, we next quantified the number of neutrophils and macrophages that were in contact with fungal cells. Throughout imaging both *A. fumigatus* and *C. albicans* infections, macrophages consistently harbored numerous phagocytosed fungi, while direct neutrophil-fungal interactions were less common (Fig. 3A-B). Quantification of the number of immune cells interacting with fungi per fish showed that macrophages were engaged with fungal cells around twice as often as neutrophils, possibly suggesting a more direct role for macrophages in controlling fungal infection in burn wounds (Fig. 3C-D).

**Figure 3.**
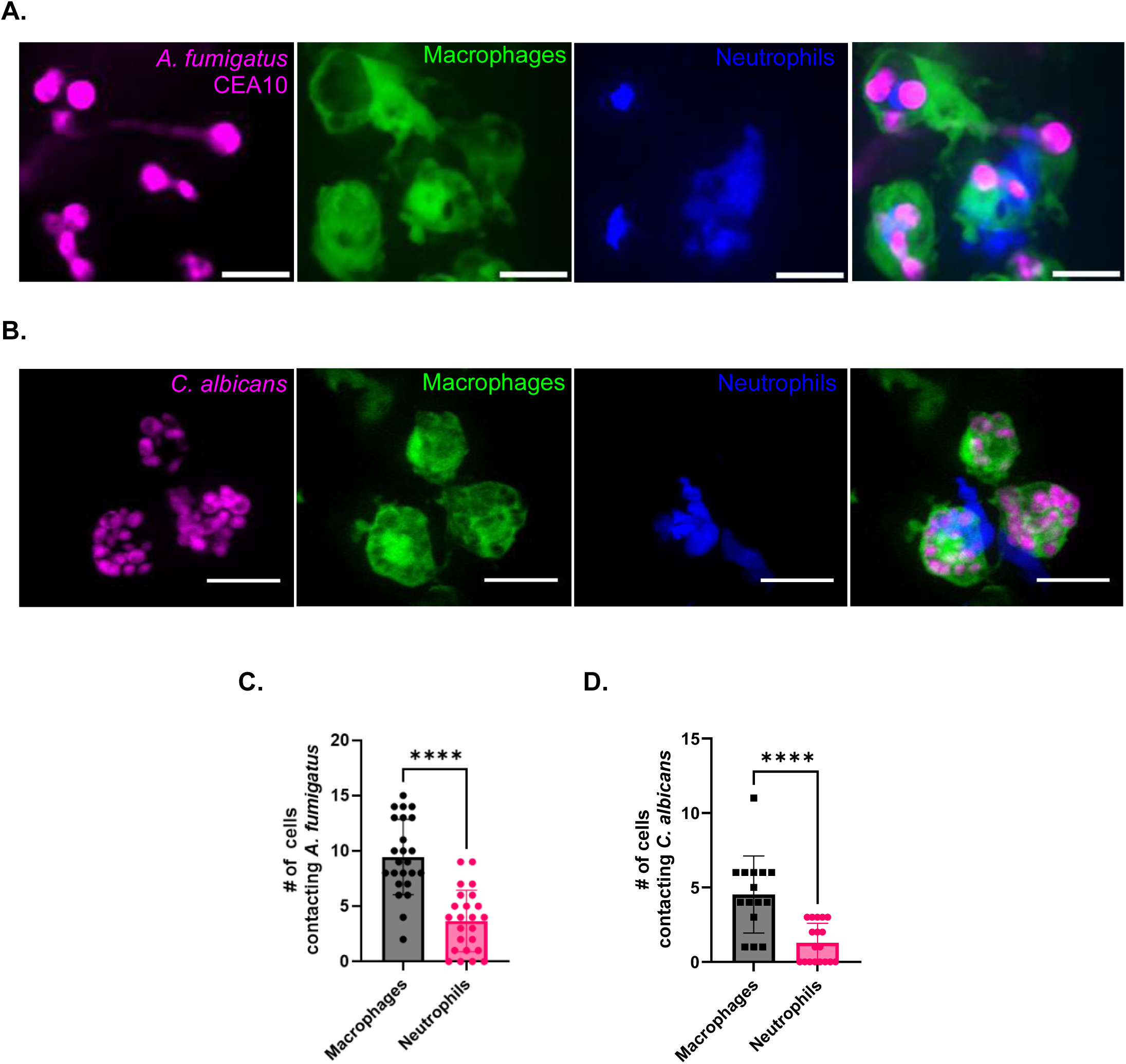
Macrophages primarily interact with fungi in burn tissue. (A-B) 24 hpb representative images of wild-type larvae with fluorescent macrophages (*mpeg-GFP*) and neutrophils (*lyz-BFP*) inoculated with fluorescent (A) *A. fumigatus* CEA10 spores (RFP) or (B) *C. albicans* yeast cells (RFP). Images represent maximum intensity projections of z-stacks (scale bar is 100µm) (C-D) Number of macrophages and neutrophils contacting (C) *A. fumigatus* cells and (D) *C. albicans* cells. Bars on graphs presented as mean + standard deviation. Each dot represents data from an individual fish and results represent data pooled from 3 independent experiments. n = 24 larvae per condition. *p* values calculated by a student’s t-test. \*\*\*\**p*<0001.

### Macrophage depletion leads to increased fungal burden in the burn wound

Our data suggest that macrophages may be the predominant leukocyte driving fungal BWI clearance (Fig. 2). However, previous work has shown that fungal phagocytosis by macrophages often does not result in fungal clearance, as both *A. fumigatus* and *C. albicans* can germinate within and escape from macrophages (21–24). To better elucidate the contribution that macrophages play in BWI resolution we transiently depleted macrophages by injecting clodronate liposomes into the caudal vein of 2 dpf larvae expressing fluorescently labeled macrophages (*mpeg-GFP*) and proceeded to burn and infect larvae at 3 dpf (25). For *A. fumigatus* infection, we found that clodronate effectively depleted macrophages and resulted in a significant increase in fungal burden at both 24 and 48 hpb when compared to PBS liposome injected larvae (Fig. 4A-C). Interestingly, we did not observe significant differences in germination, as both germ tubes and hyphae were observed throughout the experiment (Fig. 4D). However, at 72 hpb there was a slight, but insignificant, increase in germination in clodronate treated fish. Similar to *A. fumigatus,* clodronate injected fish infected with *C. albicans* showed a significant increase in fungal burden at 24 hpb (Fig. 4E-G), although this difference was lost at later time points throughout the infection process. Altogether, our data support an important role for macrophages in controlling fungal burden, especially during the first 24 hpb.

**Figure 4.**
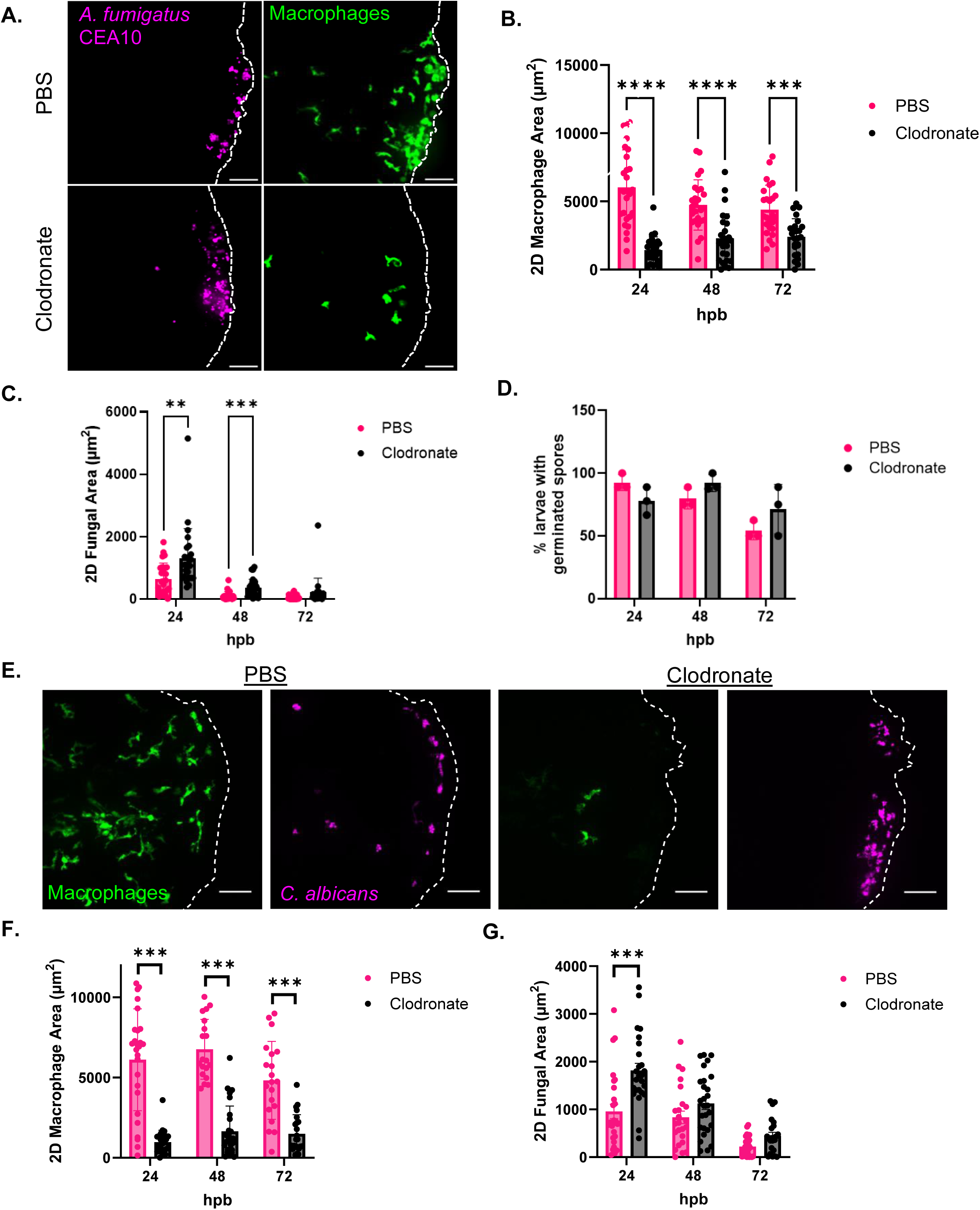
Macrophage depletion leads to increased fungal burden in the burn wound. (A) Representative images of clodronate liposome or PBS liposome treated *mpeg-GFP* larvae infected with *A. fumigatus* CEA10 at 24 hpb. Images represent maximum intensity projections of z-stacks (scale bar is 50µm). Dashed white lines outline tail fin edge. (B) 2D-macrophage area from 24-72 hpb. (C) 2D-fungal (RFP) area from 24-72 hpb. (D) Percent of larvae with germinated spores from 24-72 hpb. Each dot represents percent of an individual experiment. (E) Representative images of clodronate liposome or PBS liposome treated *mpeg-GFP* larvae infected with *C. albicans* (RFP) at 24 hpb. Images represent maximum intensity projections of z-stacks (scale bar is 50µm). Dashed white lines outline tail-fin area. (F) 2D-macrophage area from 24-72 hpb. (G) 2D-fungal (RFP) area from 24-72 hpb. Bars on graphs presented as mean + standard deviation. Each dot represents data from an individual fish and results represent data pooled from 3 independent experiments. n = 24-30 larvae per condition. *p* values calculated by a two-way ANOVA with Šidák’s multiple comparisons test. \*\**p*<0.01, \*\*\**p<*0.001, \*\*\*\**p*<0001.

### Neutrophil deficiency leads to increased hyphal growth within burn tissue

While our data place macrophages as the primary leukocyte directly interacting with fungi at the burn wound interface, there is ample evidence showing neutrophils as key players of the innate immune response against fungal infections (21, 22, 26–28). Indeed, neutrophil deficiency in human patients has been associated with invasive fungal growth and increased risk for burn wound infections, and *ex vivo* burn wound models using human tissue have shown an important role for neutrophils in the epidermal antifungal immune response (8, 29, 30). To assess the role of neutrophils during zebrafish fungal BWI, we infected larvae with impaired neutrophil function that express a dominant-negative Rac2 (RacD57N) protein under the regulatory control of a neutrophil specific promoter (*mpx:rac2D57N*). These neutrophils are not able to leave the circulation and are therefore unable to migrate to burn tissue at the infection site (31). Previous work using *mpx:rac2D57N* fish demonstrated that host-survival is decreased during *A. fumigatus* hindbrain infection, as well as for *C. albicans* swim-bladder infection (28, 32). Interestingly, we found no differences in fungal burden for *A. fumigatus* burn infection in these fish compared to WT (Fig. 5A&C). However, a significant increase in germination and hyphal development was observed in infected *mpx:rac2D57N* fish at 72 hpb (Fig. 5D). Surprisingly, a small number (3.3%) of *mpx:rac2D57N* fish presented invasive hyphal growth that extended from the wound towards the head by penetrating muscle tissue of the dorsal fin (Fig. 5B&E). Although rare, this is indicative of severe infection, in which the fungus was not only able to colonize viable tissue but to also invade beyond the infection site.

**Figure 5.**
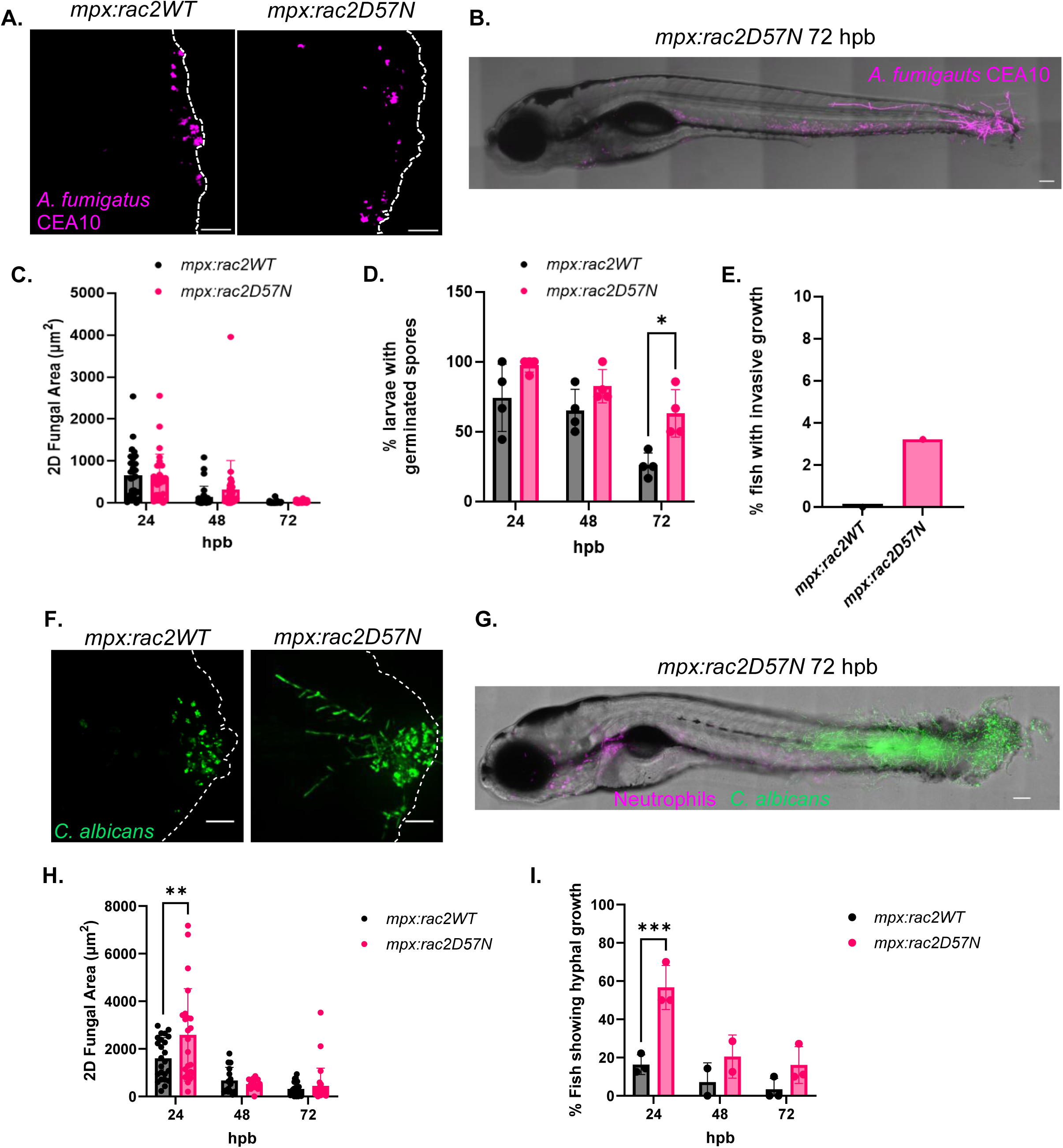
Neutrophil deficiency leads to increased hyphal growth and invasion. (A) Representative images at 24 hpb of 3-dpf *mpx*:Rac2D57N (mCherry) or *mpx*:Rac2 (mCherry) wild-type larvae infected with *A. fumigatus* CEA10 (RFP). Images represent maximum intensity projections of z-stacks (scale bar 50µm). Dashed white lines outline tail-fin area. (B) 2D-fungal (RFP) area from 24-72 hpb. (C) Percent of larvae with germinated spores from 24-72 hpb. Each dot represents percent of an individual experiment. (D) Representative image of entire *mpx*:Rac2D57N larvae infected with *A. fumigatus* CEA10 (RFP) at 72 hpb (scale bar is 200µm). (E) Percent of fish with invasive hyphal growth extending beyond the notochord and growing toward the head at 72 hpb. Bar represents the sum of fish showing invasive growth from 4 experiments. (F) Representative images at 24 hpb of 3-dpf *mpx*:Rac2D57N (mCherry) or *mpx*:Rac2 (mCherry) wild-type larvae infected with *C. albicans* (mNeon) at 24 hpb. Images represent maximum intensity projections of z-stacks (scale bar is 50µm). Dashed white lines outline tail fin edge. (G) 2D-fungal area from 24-72 hpb. (H) Representative tile image of entire *mpx*:Rac2D57N (mCherry) larvae infected with *C. albicans* (mNeon) (scale bar is 100µm). (I) Percent of fish with hyphal growth from 24-72 hpb. Each dot represents percent of an individual experiment. (Bars on graphs presented as mean + standard deviation. Each dot represents data from an individual fish and results represent data pooled from 3 independent experiments. n = 24-30 larvae per condition. *p* values calculated by a two-way ANOVA with Šidák’s multiple comparisons test. \**p*<0.05,\*\**p*<0.01, \*\*\**p<*0.001)

In contrast to *A. fumigatus*, mutant fish infected with *C. albicans* had a significant increase in fungal burden at 24 hpb compared to WT that resolved by 72 hpb (Fig. 5F and 5H). The increase in fungal burden largely correlated with extensive hyphal networks within the tail fins of *mpx:rac2D57N* infected fish by 24 hpb (Fig. 5G and 5I); ∼55% of *mpx:rac2D57N* fish showed branched hyphae extending throughout the tail fin by 24 hpb that was not observed in immunocompetent fish. Surprisingly, by 72 hpb invasive hyphal growth was absent in many of these fish, suggesting a redundant role for other immune cells in controlling fungal hyphal formation. Yet, hyphae extending beyond the notochord and growing towards the head of the zebrafish was still seen in ∼15% of the Rac2D57N infected fish (Fig. 5I), making it a more common occurrence as compared to *A. fumigatus* infection (Fig. 5E). Together, these findings highlight an essential role for neutrophils in controlling invasive hyphal growth during infection with both *A. fumigatus* and *C. albicans*.

### Impaired macrophage and neutrophil function increases invasive fungal infection

Our data have shown a role for both macrophages and neutrophils in controlling both *A. fumigatus* and *C. albicans* infections. Therefore, we hypothesized that infection in fish with deficiencies in both cell types would further exacerbate the disease phenotype. To test this, we used a zebrafish line that expresses a dominant-negative Rac2D57N in leukocytes (*coro1a:GFP-rac2D57N*) (33). Thus, both macrophages and neutrophils expressing this mutation have migration deficiencies that will impede their movement to the BWI site. Surprisingly, infection of *coro1a:GFP-rac2D57N* fish with *A. fumigatus* did not appear to result in increased fungal burden or germination rates (Fig. 6A-C), but did result in an increase in the number of fish showing invasive hyphal growth extending beyond the notochord as early as 24 hpb. Indeed, 10% and 5% of all fish imaged showed hyphal growth extending past the notochord and moving toward the head at 24 and 48 hpb, respectively (Fig. 6D). Therefore, loss of both macrophages and neutrophils further impacts invasive hyphal growth of *A. fumigatus*, although it appears to be modest.

**Figure 6.**
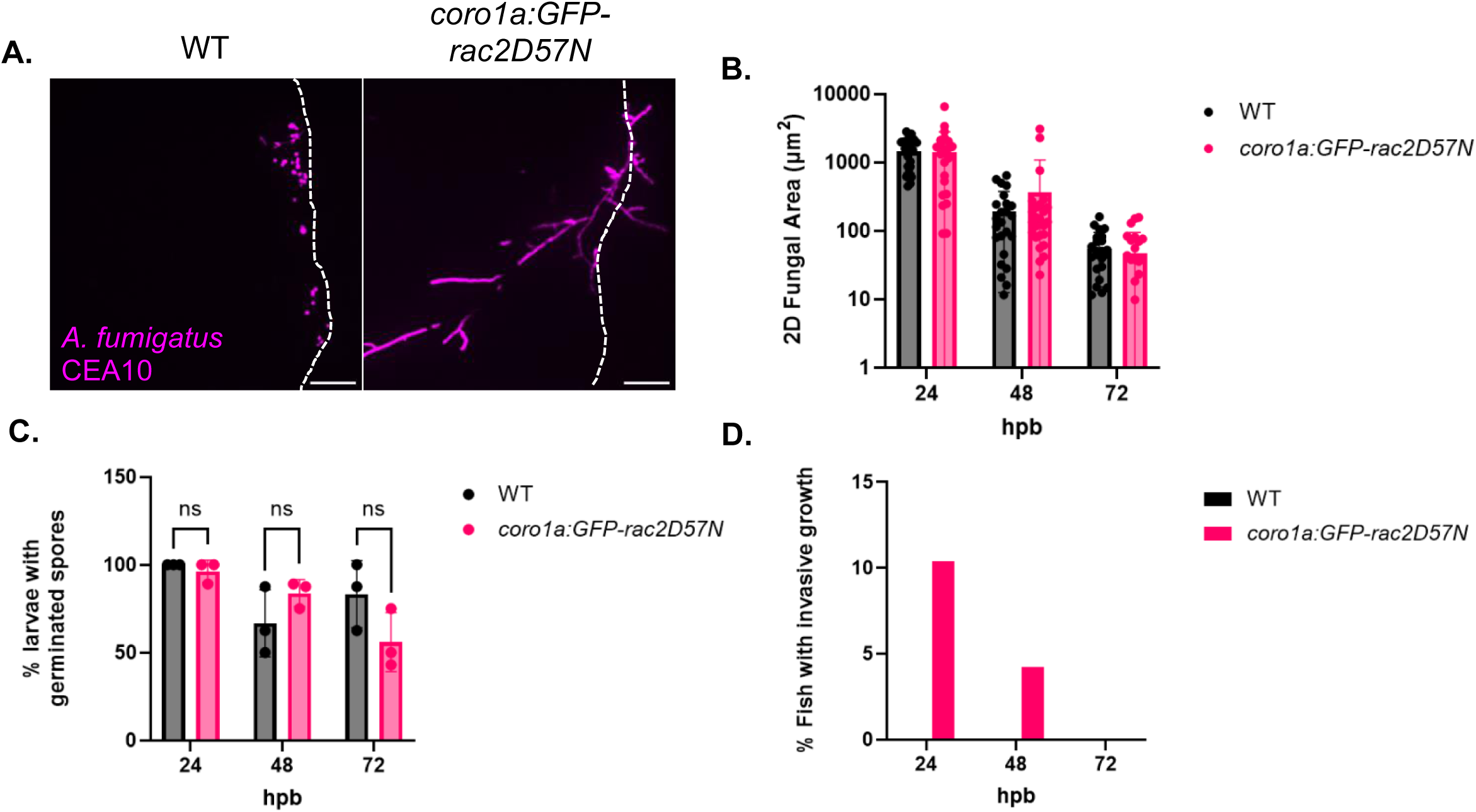
Impaired macrophage and neutrophil function results in increased *A. fumigatus* invasive hyphal growth. (A) Representative images of 3-dpf *coro1a:GFP-rac2D57N* (GFP) and wild-type larvae infected with *A. fumigatus* CEA10 (RFP) at 24 hpb. Images represent maximum intensity projections of z-stacks (scale bar is 50µm). (B) 2D-fungal area (RFP) at 24-72 hpb. (C) Percent of larvae with germinated spores at 24-72 hpb. Each dot represents percent of an individual experiment. (B) Percent of fish with invasive hyphal growth at 72 hpb. Bar represents the sum of fish with invasive hyphal growth from 3 experiments. Bars on graphs presented as mean + standard deviation. Each dot represents data from an individual fish and results represent data pooled from 3 independent experiments. n = 24-30 larvae per condition. *p* values calculated by a two-way ANOVA with Šidák’s multiple comparisons test.

Unlike *A. fumigatus*, infection in *coro1a:GFP-rac2D57N* fish with *C. albicans* showed a significant increase in fungal burden at all the times analyzed (Fig. 7A-C). This was accompanied by robust hyphal development as early as 24 hpb that persisted throughout infection. Overall, between 50-75% of *C. albicans* infected *coro1a:GFP-rac2D57N* zebrafish showed hyphal growth of varying degrees throughout the infection (Fig. 7D). We then further categorized hyphal presence based on the severity of invasive growth (Fig. 7E). At 24 hpb, the hyphal growth was either small hyphae localized to the burn wound edge, denoted as level 1, or extensive and branched hyphal growth that was localized within the tail fin, denoted as level 2 (Fig. 7F). However, as the infection progressed the percentage of fish displaying hyphae extending past the notochord and towards the head of the fish, denoted as very invasive level 3 hyphal growth, increased. Indeed, ∼55% and ∼80% of hyphal growth at 48 hpb and 72 hpb, respectively, were classified as severely invasive (Fig. 7F). Therefore, it appears that both macrophages and neutrophils are essential to control *C. albicans* during infection. However, it is important to note that *coro1a:GFP-rac2D57N* fish also showed an innate sensitivity to burn wound injury, and excessive tissue necrosis and death were observed even in the absence of infection (Supplementary Fig. 2A-B). Fungal presence did not appear to further increase death, and it seems likely that the excessive tissue damage observed in *coro1a:GFP-rac2D57N* fish allowed *C. albicans* to more effectively colonize and invade the burn wound niche.

**Figure 7:**
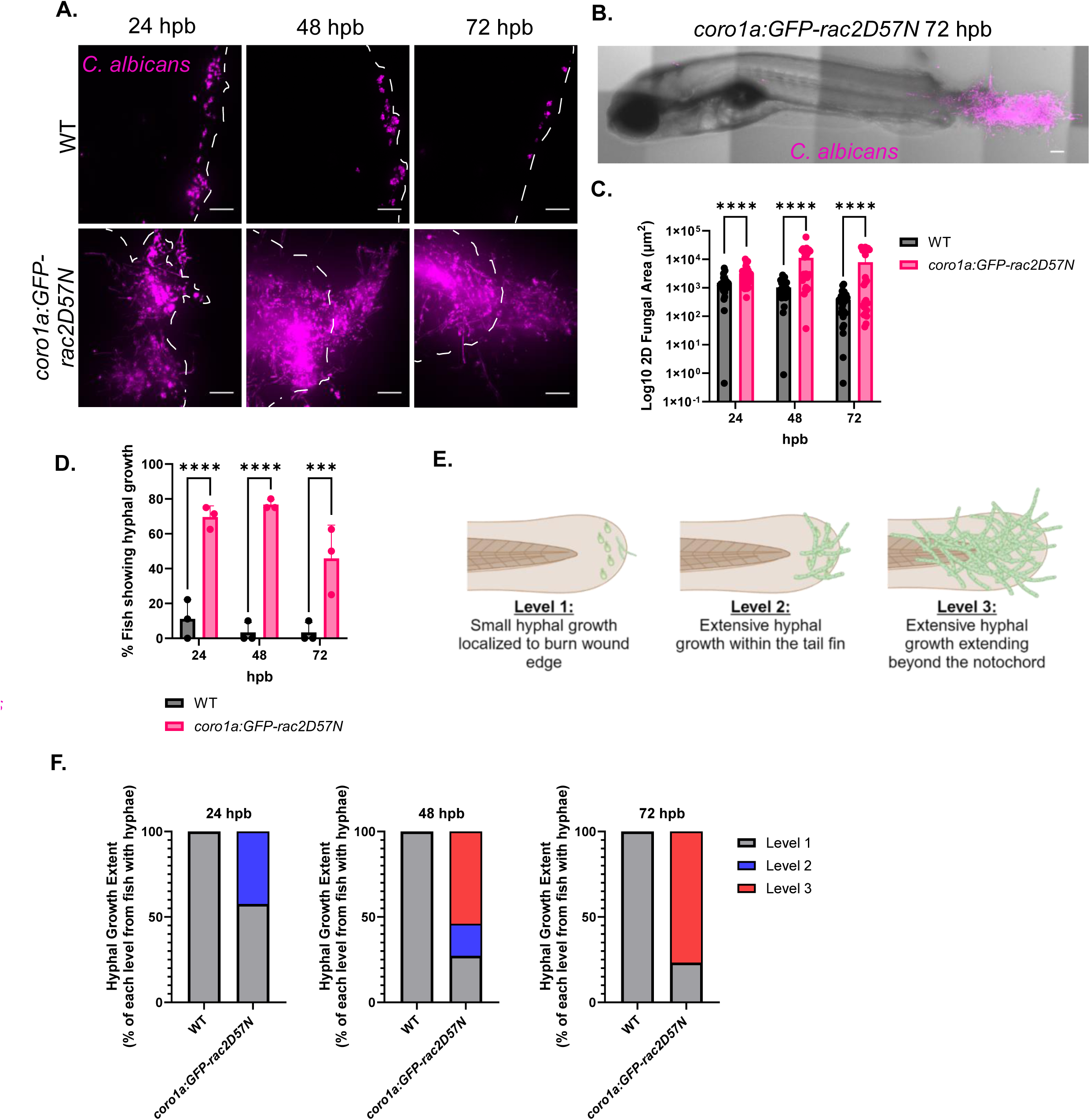
Impaired macrophage and neutrophil function results in uncontrolled growth and invasion of *C. albicans* in thermal injury. (A) Representative images of 3-dpf *coro1a:GFP-rac2D57N* (GFP) or wild-type larvae infected with *C. albicans* (dTomato) and imaged at 24, 48 and 72-hpb. Images represent maximum intensity projections of z-stacks (scale bar is 50µm). Dashed white lines outline tail fin edge. (B) Representative tile images of entire *coro1a:GFP-rac2D57N* (GFP) larvae infected with *C. albicans* (dTomato) at 72 hpb. (scale bar is 100µm). (C) Log_10_ 2D-fungal area from 24-72 hpb. (D) Percent of fish showing hyphal growth from 24-72 hpb. Each dot represents percent of an individual experiment. (D) Schematic of hyphal growth levels classification. Level 1 indicates small and controlled hyphal growth present at the burn wound edge. Level 2 is denoted as extensive hyphal growth within the tail fin. Level 3 is classified as extensive and invasive hyphal growth that penetrates beyond the notochord and extends towards the head of the fish. (F) Percent of fish showing hyphal growth within each classification level (1-3). Bar colors represent the different levels and are derived from the mean of 3 individual experiments. (Bars on graphs presented as mean + standard deviation. Each dot represents data from an individual fish and results represent data pooled from 3 independent experiments. n = 24-30 larvae per condition. *p* values calculated by a two-way ANOVA with Šidák’s multiple comparisons test. \*\*\**p<*0.001, \*\*\*\**p*<0001)

### Increased β(1,3)-glucan exposure attenuates *C. albicans* burn wound colonization in zebrafish

Both macrophages and neutrophils are needed to control burn wound infection by *C. albicans* (Fig. 7) and we hypothesized that infection with fungal mutants that have an attenuated ability to evade host immune system recognition would result in enhanced fungal clearance within this niche. *C. albicans* mutants with a hyperactive Cek1 mitogen activated protein kinase pathway (*STE11/P_tet-off_STE11^ΔN467^*) or that are deficient in the cell wall protein Fgr41 (*fgr41 Δ/Δ*) display increased exposure of the immunogenic cell wall epitope β(1,3)-glucan (Fig. 8A), induce increased TNFα production by macrophages *in vitro* and display virulence attenuations that are host immune system dependent during systemic infection in mice (34–36). Therefore, we predicted that these mutants would also be attenuated in their ability to colonize burn tissue within an immunocompetent zebrafish burn model. Indeed, WT fish infected with either *STE11/P_tet-off_STE11^ΔN467^* or *fgr41Δ/Δ* showed significantly less fungal burden than WT *C. albicans* infected fish at both 24 hpb and 48 hpb (Fig. 8B). Thus, masking immunogenic β(1,3)-glucan is an important mechanism utilized by *C. albicans* to successfully colonize burn tissue.

**Figure 8.**
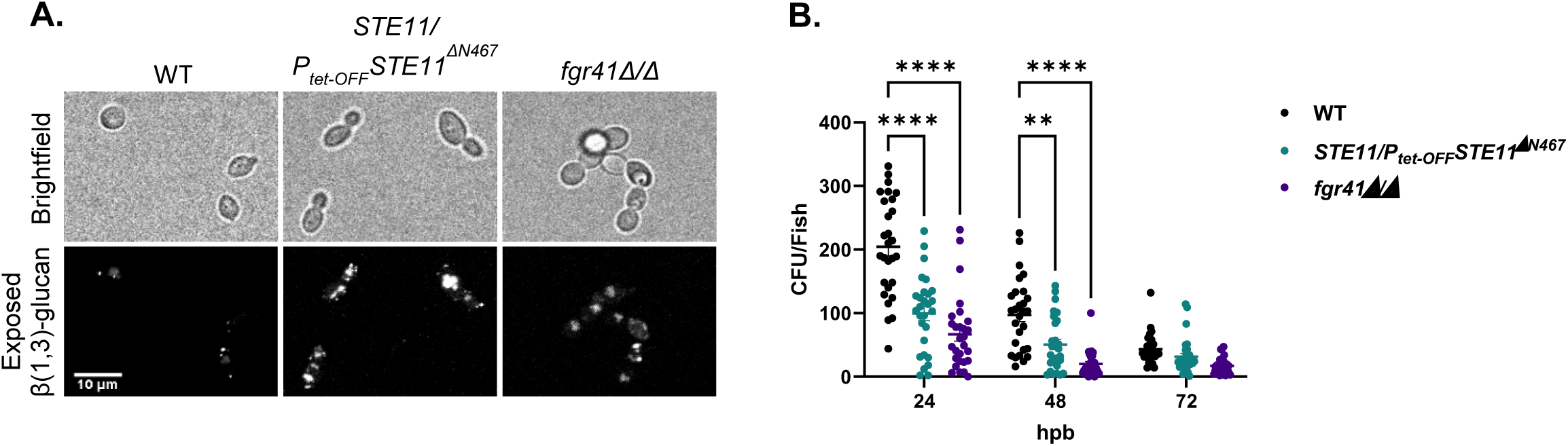
Increased β-glucan exposure attenuates burn wound colonization in zebrafish. (A) Representative microscopy images of β(1,3)-glucan exposure of wild-type, *STE11/P_tet-OFF_STE11^ΔN467^* and *fgr41Δ/Δ C. albicans* strains. The scale bar indicates 10μm. (B) CFUs per fish from wild-type zebrafish larvae that were infected with either the *C. albicans* wild-type or mutants that show increased β(1,3)-glucan exposure (*STE11/P_tet-OFF_STE11^ΔN467^* and *fgr41Δ/Δ*). Error bars on graph presented as mean + standard deviation. Each dot represents data from an individual fish. Results represent data pooled from 3 independent experiments. n= 24-30 larvae per condition. *p* values calculated by a two-way ANOVA with Šidák’s multiple comparisons test. \*\**p*<0.01, \*\*\*\**p*<0001.

## Discussion

Fungal burn wound infections are an understudied, but significant complication that increases the risk of disease and mortality in patients. The challenge of finding suitable animal models to study the biology behind fungal infections during burn injury represents a primary obstacle to better understanding disease kinetics. Because of this, most of the available data is limited to clinical studies, *ex vivo* models and sparse *in vivo* studies that have primarily focused on reactive treatment strategies after infection (8, 10, 12, 37, 38). Therefore, there remain significant gaps in our understanding of the basic biology of the fungal-host burn wound interface. In this study we exploited an already established burn injury model in zebrafish larvae and developed a fungal burn wound infection model using two of the most common species isolated from burn wounds, *A. fumigatus* and *C. albicans*.

By combining live imaging and quantification of fungal burden, we demonstrate that both *A. fumigatus* and *C. albicans* successfully colonize and are cleared from immunocompetent zebrafish larvae (Fig. 1). Both macrophages and neutrophils are recruited to the site of trauma (Fig. 2). Fungal colonization results in an increase in recruitment of neutrophils in response to both pathogens, but only *C. albicans* induces increased macrophage infiltration as well. Thus, it appears that each fungal isolate has distinct effects on the host immune response following burn injury. It is possible that the divergent metabolic nature between the pathogens may account for these differences. *C. albicans* yeast cells are active and appear to readily undergo morphological transitioning to hyphae at the early stages (2-6 hpb) of burn wound colonization (Fig. 1). Hyphae are both more invasive than yeast cells, but also have increased exposure of β-glucan moieties within their cell wall that may further exacerbate the host immune response (39–41). By contrast, *A. fumigatus* conidia are generally considered metabolically dormant, and germination does not occur until 24-48 hpb (Fig. 1) (42). Structurally, the spore surface is covered by hydrophobins that mask underlying immunogenic cell wall epitopes and aids in immune cell evasion (43, 44). Therefore, the delayed germination rate of *A. fumigatus* spores likely contributes to the reduced immune cell infiltration as compared to *C. albicans*.

Our findings suggest distinct roles for neutrophils and macrophages at different stages of infection. Depletion of macrophages results in increased fungal burden at early stages of infection (Fig. 4), while infection in neutrophil defective zebrafish results in invasive hyphal growth by both fungi (Fig. 5). However, both fungal burden and hyphal presence are largely restored to WT levels at the later stages of infection in both macrophage or neutrophil deficient zebrafish, suggesting redundant functions. Infection in *coro1a:GFP-rac2D57N* zebrafish larvae, which have defective neutrophils and macrophages, increases both fungal burden and invasive hyphal growth with *C. albicans* (Fig. 7), supporting the idea that macrophages and neutrophils have synergistic and partially redundant roles in antifungal immunity in burns. The observation that infection in fish with defective neutrophils results in increased invasive hyphal development (Fig. 5) recapitulates findings from both clinical reports and *ex vivo* human skin infection models (8, 30). However, our findings that loss of macrophages impacts fungal colonization (Fig. 4) raises interesting questions about the role of macrophages in fungal control during thermal injury. *C. albicans* yeast cells can remain viable, divide and be transported within macrophages to distant sites from the initial infection (45). Studies with *A. fumigatus* using the zebrafish hindbrain infection model have similarly shown that macrophages phagocytose but often do not kill conidia, and actually delay clearance by inhibiting germination and subsequent hyphal killing by neutrophils (22). Yet, it was recently reported that macrophage specific Rac2 rescues control of invasive hyphal growth by *A. fumigatus* in pan-Rac2 deficient fish during hindbrain infection (46). Our data further indicates that macrophages can control hyphal growth in the absence of neutrophils, demonstrating that macrophages do play an important role in antifungal immunity. A caveat is that there may be off target effects of clodronate treatment that contribute to some of the observed phenotypes, since studies in mice suggest that clodronate liposomes may also have effects on neutrophil function (47).

Our findings highlight a distinction between colonization and infection. In humans, skin infection is characterized by fungal penetration to variable depths within viable tissue, along with angioinvasion (48–50). Colonization is limited to the wound surface and proliferation at the interface between viable and non-viable tissue (9, 49, 51). In immunocompetent and macrophage depleted fish, fungi were highly localized and displayed minimal hyphal development, which suggests colonization of the tissue. However, neutrophil deficiency was associated with fungal invasion of neighboring tissues. Although zebrafish larvae do not fully develop vasculature until later stages of development, we propose that the invasive hyphal growth observed in larvae with impaired neutrophil function represents infected tissue and supports a key role of neutrophils in preventing invasive growth within burns (52).

To test the utility of this fungal burn wound model for identifying fungal mechanisms of tissue colonization, we infected fish with *C. albicans* mutants that had increased β(1,3)-glucan exposure within their cell walls (Fig. 8). β(1,3)-glucan is highly immunogenic, and these mutants have been previously shown to induce robust pro-inflammatory cytokine production by macrophages *in vitro* and show immune system-dependent virulence attenuations during systemic infections in mice (34–36). Our results show limited tail fin colonization with unmasked *C. albicans* mutants (Fig. 8), similar to the phenotypes observed in mice. These findings suggest that masking of β(1,3)-glucan is necessary for persistent tissue colonization in larval burn wounds, and highlights the efficacy of this model for studying fungal mechanisms of tissue colonization during burn injury. However, further analysis should be performed to better characterize the host response to infection with cell wall mutants within the context of BWIs, with the potential for these findings to have broader implications across different infection models and disease manifestations.

## Materials and Methods

### Ethics Statement

The use of zebrafish in this research was approved by the Institutional Animal Care and Use Committee (IACUC) at University of Wisconsin-Madison College of Agricultural and Life Sciences (CALS). The animal care and use protocol M005405-A02 adheres to the guidelines established by the federal Health Research Extension Act and the Public Health Service Policy on the Humane Care and Use of Laboratory Animals, led by the National Institutes of Health (NIH) Office of Laboratory Animal Welfare (OLAW).

### Zebrafish Husbandry and Maintenance

Adult zebrafish were maintained under a light/dark cycle of 14/10 hours, as previously described (32). For experiments, embryos were collected, screened and maintained at 28.5°C. For *A. fumigatus* infection in wild-type Rac2, zebrafish larvae that weren’t expressing mCherry were chosen as the fungus was RFP positive. All zebrafish lines utilized in this study are listed in Table 1.

| Line | Description | Reference |
| --- | --- | --- |
| WT (AB) | Wild-type line | ZIRC |
| Tg(lyz:BFP/mpeg1:GFP) | Wild type with BFP-expressing neutrophils and GFP-expressing macrophages | (22) |
| Tg(mpeg1:GFP) | Wild type with GFP-expressing macrophages | ZIRC |
| Tg(lyz:GFP) | Wild type with GFP-expressing neutrophils | ZIRC |
| Tg(mpx:mCherry-2A-rac2D57N) | Rac2D57N mutation and mCherry-expressing neutrophils | (31) |
| Tg(mpx:mCherry-2A-rac2WT) | Wild-type Rac2 and mCherry-expressing neutrophils | (31) |
| Tg(*coro1a*:GFP-rac2D57N)<sup>psi92Tg</sup> | Rac2D57N mutation and GFP expressing macrophages and neutrophils | (33) |

### Fungal strains growth and conditions

*Aspergillus fumigatus* and *C. albicans* strains were maintained as glycerol stocks at −80°C. For *A. fumigatus,* the strain was activated before every experiment by streaking on a plate of Glucose Minimal Media (GMM) and incubating for 3 days in darkness at 37°C. Conidia were harvested in 0.01% Tween-water by scraping with an L-shaped spreader, and immediately filtered through sterile Miracloth. To prepare for burn wound infection, conidia solution was centrifuged and washed three times in sterile 1x DPBS (-CaCl_2_, -MgCl_2_). After counting conidia, solution was resuspended in PBS. The appropriate volume to achieve 2 x 10^7^ spores/mL was then added to a conical tube (per dish of larvae) with 5 mL of E3 without methylene blue (-MB).

For *C. albicans,* strains were activated on YPD agar plates (1% Yeast Extract, 2% peptone, 2% dextrose, 2% agar) from glycerol stocks. One day prior to infection, overnight cultures in liquid YPD media were started and left to incubate for ∼16 hours while shaking at 225rpm at 30°C. Following incubation, cultures were transferred to a 15ml conical tube and centrifuged at 3500rpm for 5 minutes. Following centrifugation, the supernatant was aspirated, and the fungal cell pellet was washed 2 times with 10ml of 1x DPBS. After washing, the fungal cell pellet was resuspended in 5ml of E3 -MB media, counted on a hemocytometer, and diluted to a concentration of 2×10^7^ cells/ml in 6ml of E3 -MB media. All fungal strains used in this study are listed in Table 2.

For fixation of fungal cells, *A. fumigatus* and *C. albicans* cells were incubated in 4% Paraformaldehyde (PFA) for 30 minutes at room temperature and washed three times in 1x DPBS.

| Strain | Parent | Genotype | Reference |
| --- | --- | --- | --- |
| *Aspergillus fumigatus* |  |  |  |
| TDGC1.2 | TCDN6.7 | $\Delta akuB$; $argB^-$; $gpdA::RFP::argB$; $pyrG^-$; $fumipyrG$ | (53) |
| *C. albicans* |  |  |  |
| Caf2-dTomato | Caf2 | $\Delta ura3::imm434/URA3 pENO1-dTomato-NATR$ | (54) |
| SC5314-mNeon | SC5314 | $P_{ENO1}-NEON-NAT$ (Wild Type) | (55) |
| AWY006 | SC5314 | $LEU2/leu2\Delta$ (Wild Type) | (36) |
| AWY080 | AWY006 | $STE11/P_{tet-OFF}-STE11^{\Delta N467} LEU2/leu2\Delta$ | (36) |
| SLY005 | AWY006 | $fgr41\Delta/\Delta LEU2/leu2\Delta$ | (35) |

### Thermal injury and infection

Before every experiment, larvae were anesthetized in E3 -MB containing 0.2 mg/mL Tricaine (ethyl 3-aminobenzoate, Sigma). For thermal injury and infection, larvae were placed in 10 mL of E3 -MB + Tricaine in a 60 mm tissue culture treated dish (8-10 larvae per dish). A fine tip cautery pen (Bovie, Symmetry Surgical, Antioch, TN) was used to burn the caudal fin without injuring the notochord (16). Immediately after burn, 5 mL of liquid was removed from the dish and replaced with 5 mL of fungal solution (containing 2×10^7^ cells/ml of fungi) to achieve a final concentration of 1 x 10^7^ cells/mL. For control larvae, this was done with 5 mL of E3 -MB + 100uL of PBS. Larvae were slowly shaken and kept in solution for 1 hour. Afterwards, larvae were rinsed five times with E3 -MB and incubated at 28.5°C for further analysis.

### Clodronate liposome injection

Transgenic larvae with fluorescent macrophages (*mpeg: GFP*) were microinjected at 2dpf into the caudal vein plexus with 2nL of clodronate or PBS liposomes (Liposoma) with 0.1% phenol red for easier visualization (25).

### Fungal Burden Enumeration by colony forming units

For *A. fumigatus* and *C. albicans*, individual larvae were collected at 24, 48 and 72 hpb and placed in 1.5 ml microcentrifuge tubes with 90uL of 1x DPBS with 50ug/mL gentamycin and 50ug/mL kanamycin. Larvae were homogenized for 15 seconds in a mini bead beater at maximum speed. The entire volume was plated on solid GMM (*A. fumigatus*) or YPD (*C. albicans*) and plates were incubated at 30-37°C for 2 days and CFUs were counted. Three biological replicates were run for all cell lines tested and the total number of fish used is listed in the appropriate figure legends. Statistical analysis was performed using a one-way ANOVA with a Tukey’s multiple comparison analysis (GraphPad Prism, v7.0c software).

### Live imaging acquisition

For live imaging acquisition, larvae were anesthetized and embedded flat on their side in a 35mm glass bottom dish (CellVis) by pouring 1-2% low melting point agarose (E3 -MB) on top. Z-series images (5µm slices) of the tail fin were acquired on a spinning disk confocal microscope (CSU-X; Yokogawa) with a confocal scan head on a Zeiss Observer Z.1 inverted microscope, Plan-Apochromat NA 0.8/20x objective, and a Photometrics Evolve EMCCD camera. All images were acquired using ZEN 2.6 software. Tiles and Z-series were acquired and stitched using ZEN software for whole larvae images.

### Imaging analyses and processing

Images were processed and analyzed using FIJI ImageJ (56). Maximum intensity projections of Z-series images were used for representative images and for further analysis. Fungal burden and leukocyte recruitment were analyzed by manual thresholding and measuring 2D area of the corresponding fluorescent signal. Germination for *A. fumigatus* was scored as the presence or absence of germinated conidia (germ tube or hyphae). Invasive hyphal growth was quantified by the presence of branched hyphal networks extending beyond the tail fin edge toward the caudal vein of the fish. For immune cells contacting fungus analysis, each image in the whole Z-series was analyzed and the number of macrophages or neutrophils physically contacting fungal cells within the same image plane were counted. Counts were then pooled for all planes to generate contact numbers per fish. In all representative images, brightness and contrast histograms were adjusted for visual purposes only, and no alterations were made to images prior to analysis. For measuring tissue regrowth, the total fin tissue area distal to the notochord was outlined using the polygon tool (16). Three biological replicates were run for all experiments and the total number of fish used is listed in the appropriate figure legends. Statistical analysis was performed using a two-way ANOVA with a Šidák’s multiple comparisons test (GraphPad Prism, v7.0c software).

### Β(1,3)-glucan staining

Fungal Β(1,3)-glucan was stained as previously described, with modification to the secondary antibody used in this study (57). Here, a secondary rat-anti-mouse FITC conjugated antibody (Biolegend; 406001) was used at a 1:250 dilution for staining. Following staining, cells were imaged on a Zeiss Observer Z.1 inverted microscope and images processed via Fiji ImageJ.

## Acknowledgements

We would like to thank Dr. Robert Wheeler (University of Maine) for providing the fluorescent *C. albicans* strains used in this study. Burn wound infection schematic (Figure 1) created in BioRender. Huttenlocher, A. (2024) https://BioRender.com/i26e377

---

**Supplementary figure 1.**
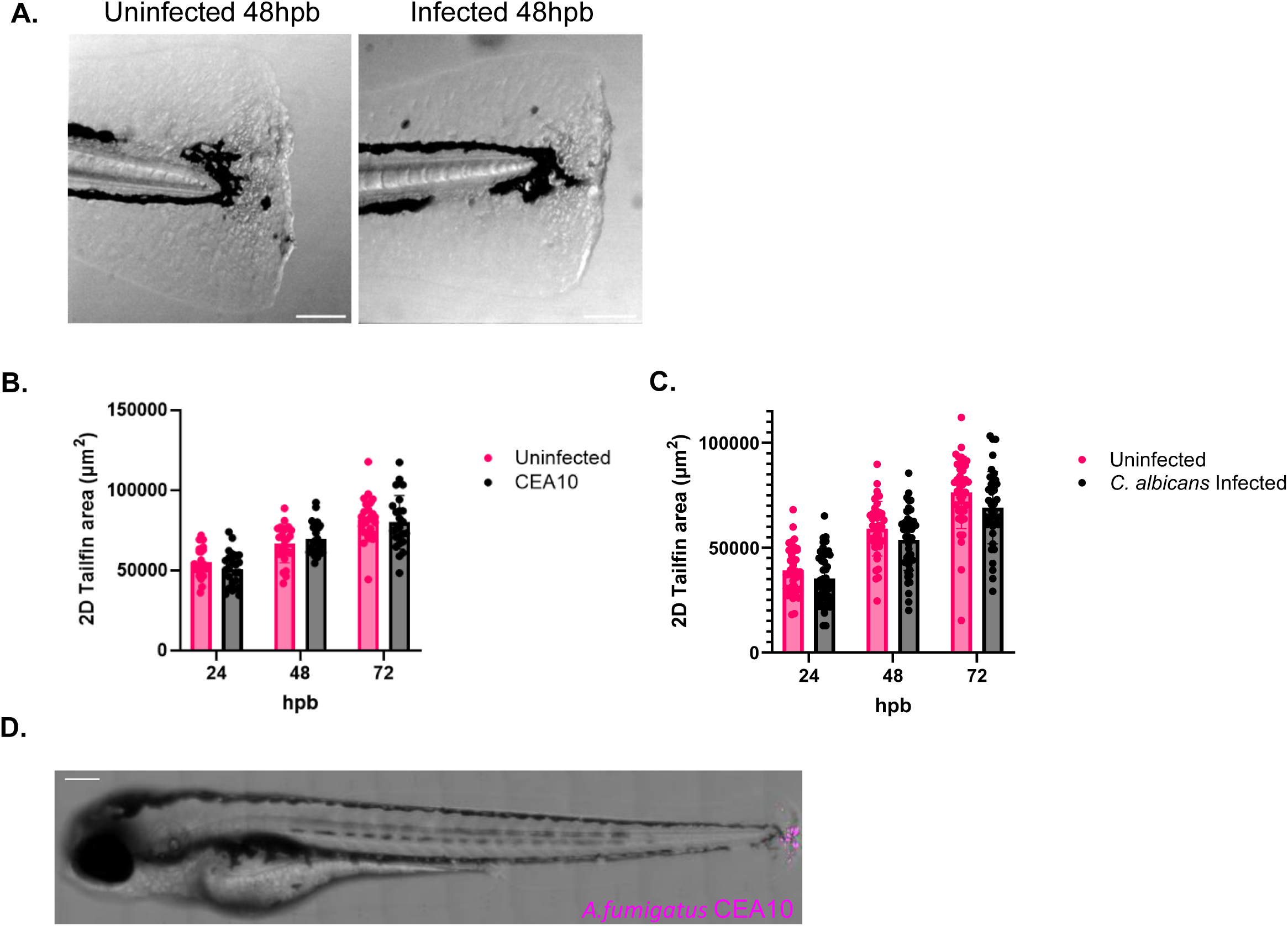
Tail fin regrowth is not impaired by infection and infection is localized to the tail region. 3-dpf wild-type larvae were injured and infected with either *A. fumigatus* CEA10 (RFP) or *C. albicans* (RFP). Control fish were uninfected. (A) Representative brightfield images of larvae infected with *A. fumigatus* CEA10 or uninfected at 48hpb. (B) 2D-tail fin area at 24-72 hpb of larvae uninfected or infected with *A. fumigatus* CEA10. (C) 2D-tail fin area at 24-72 hpb of larvae uninfected or infected with *C. albicans.* (D) Representative tile image of entire wild-type larvae infected with *A. fumigatus* CEA10 (RFP) (scale bar 200 µm). Bars on graph presented as mean+s.d. Each dot represents data from an individual fish. Results represent data pooled from 3 independent experiments. n= 24-30 larvae per condition. *p* values calculated by ANOVA with Tukey’s multiple comparisons

**Supplementary Figure 2.**
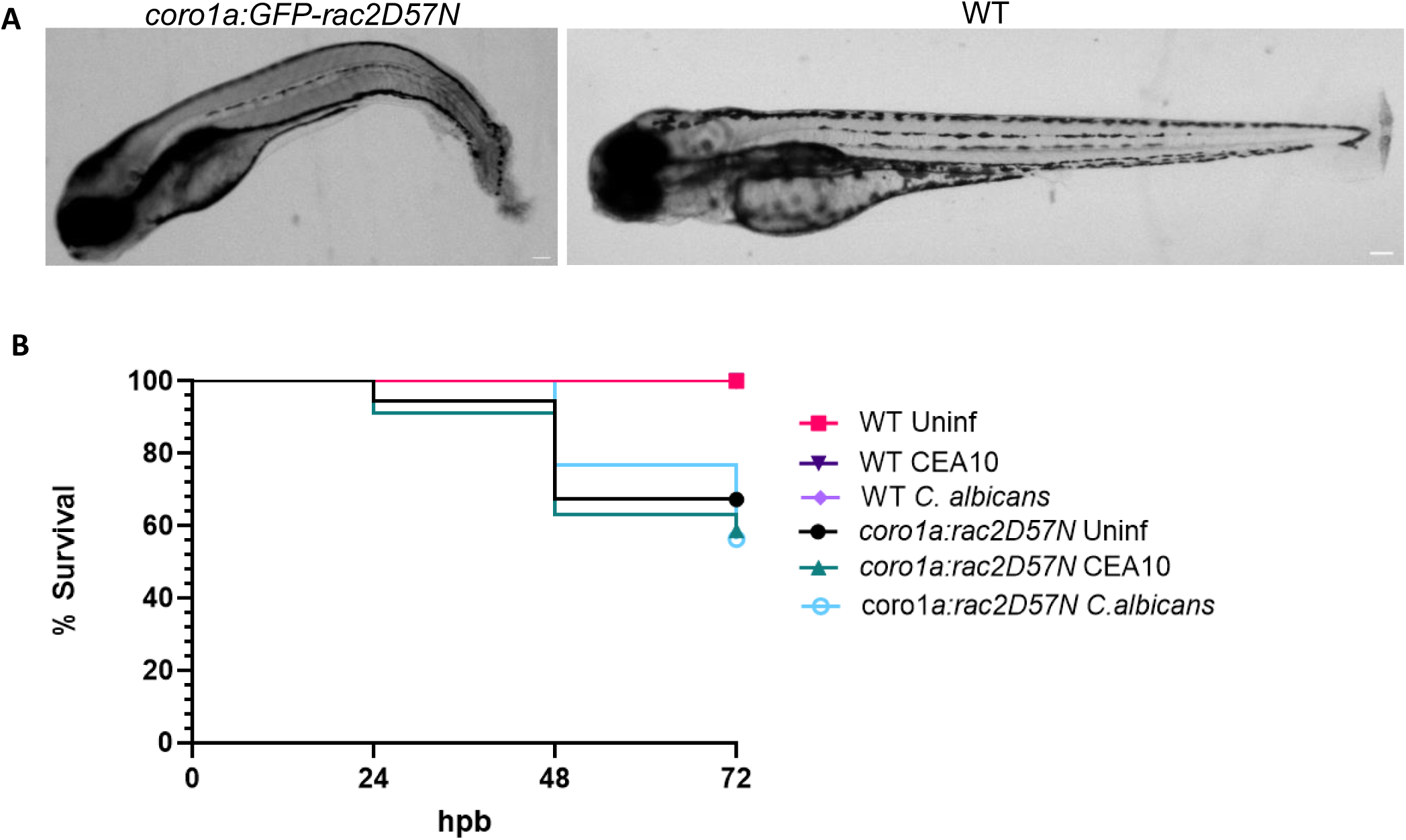
Macrophage and neutrophil deficient larvae are more susceptible to burn wound. 3-dpf *coro1a:GFP-rac2D57N* (GFP) or wild-type larvae were injured and infected with *A. fumigatus* CEA10 or *C. albicans* and imaged at 24 hpb. (A) Representative brightfield tile image of entire uninfected wild-type or *coro1a*:Rac2D57N larvae infected (scale bar 200µm). (B) Fish were monitored for survival at 24-72hpb. Survival curve of infected versus uninfected wild-type and *coro1a:GFP-rac2D57N* larvae.

## References

1. Jeschke MG, van Baar ME, Choudhry MA, Chung KK, Gibran NS, Logsetty S. 2020. Burn injury. Nat Rev Dis Primers 6:11.

2. Church D, Elsayed S, Reid O, Winston B, Lindsay R. 2006. Burn Wound Infections. Clin Microbiol Rev 19:403–434.

3. Roy S, Mukherjee P, Kundu S, Majumder D, Raychaudhuri V, Choudhury L. 2024. Microbial infections in burn patients. Acute Crit Care 39:214–225.

4. Pruskowski KA, Mitchell TA, Kiley JL, Wellington T, Britton GW, Cancio LC. 2021. Diagnosis and Management of Invasive Fungal Wound Infections in Burn Patients. 4. European Burn Journal 2:168–183.

5. Capoor MR, Sarabahi S, Tiwari VK, Narayanan RP. 2010. Fungal infections in burns: Diagnosis and management. Indian J Plast Surg 43:S37–S42.

6. Branski LK, Al-Mousawi A, Rivero H, Jeschke MG, Sanford AP, Herndon DN. 2009. Emerging Infections in Burns. Surg Infect (Larchmt) 10:389–397.

7. Ballard J, Edelman L, Saffle J, Sheridan R, Kagan R, Bracco D, Cancio L, Cairns B, Baker R, Fillari P, Wibbenmeyer L, Voight D, Palmieri T, Greenhalgh D, Kemalyan N, Caruso D. 2008. Positive Fungal Cultures in Burn Patients: A Multicenter Review: Journal of Burn Care & Research 29:213–221.

8. von Müller C, Bulman F, Wagner L, Rosenberger D, Marolda A, Kurzai O, Eißmann P, Jacobsen ID, Perner B, Hemmerich P, Vylkova S. 2020. Active neutrophil responses counteract Candida albicans burn wound infection of ex vivo human skin explants. Sci Rep 10:21818.

9. Struck MF, Gille J. 2013. Fungal infections in burns: a comprehensive review. Ann Burns Fire Disasters 26:147–153.

10. Abdullahi A, Amini-Nik S, Jeschke MG. 2014. Animal models in burn research. Cell Mol Life Sci 71:3241–3255.

11. Maslova E, Osman S, McCarthy RR. 2023. Using the Galleria mellonella burn wound and infection model to identify and characterize potential wound probiotics. Microbiology (Reading) 169:001350.

12. Maslova E, Shi Y, Sjöberg F, Azevedo HS, Wareham DW, McCarthy RR. 2020. An Invertebrate Burn Wound Model That Recapitulates the Hallmarks of Burn Trauma and Infection Seen in Mammalian Models. Front Microbiol 11:998.

13. Rosowski EE. 2020. Determining macrophage versus neutrophil contributions to innate immunity using larval zebrafish. Disease Models & Mechanisms 13:dmm041889.

14. Rosowski E, Knox B, Archambault L, Huttenlocher A, Keller N, Wheeler R, Davis J. 2018. The Zebrafish as a Model Host for Invasive Fungal Infections. JoF 4:136.

15. Robertson TF, Huttenlocher A. 2022. Real-time imaging of inflammation and its resolution: It’s apparent because it’s transparent*. Immunological Reviews 306:258–270.

16. Miskolci V, Squirrell J, Rindy J, Vincent W, Sauer JD, Gibson A, Eliceiri KW, Huttenlocher A. 2019. Distinct inflammatory and wound healing responses to complex caudal fin injuries of larval zebrafish. eLife 8:e45976.

17. Fister AM, Horn A, Lasarev MR, Huttenlocher A. 2024. Damage-induced basal epithelial cell migration modulates the spatial organization of redox signaling and sensory neuron regeneration. eLife 13:RP94995.

18. Barros-Becker F, Squirrell JM, Burke R, Chini J, Rindy J, Karim A, Eliceiri KW, Gibson A, Huttenlocher A. 2020. Distinct Tissue Damage and Microbial Cues Drive Neutrophil and Macrophage Recruitment to Thermal Injury. iScience 23:101699.

19. Horn A, Wagner AS, Hou Y, Zajac JC, Fister AM, Chen Z, Pashaj J, Junak M, Soto NMM, Gibson A, Huttenlocher A. 2024. Isotonic medium treatment limits burn wound microbial colonization and improves tissue repair. bioRxiv 10.1101/2024.10.29.620892.

20. Capoor MR, Gupta S, Sarabahi S, Mishra A, Tiwari VK, Aggarwal P. 2012. Epidemiological and clinico-mycological profile of fungal wound infection from largest burn centre in Asia. Mycoses 55:181–188.

21. Brothers KM, Newman ZR, Wheeler RT. 2011. Live Imaging of Disseminated Candidiasis in Zebrafish Reveals Role of Phagocyte Oxidase in Limiting Filamentous Growth. Eukaryotic Cell 10:932–944.

22. Rosowski EE, Raffa N, Knox BP, Golenberg N, Keller NP, Huttenlocher A. 2018. Macrophages inhibit Aspergillus fumigatus germination and neutrophil-mediated fungal killing. PLoS Pathog 14:e1007229.

23. Jia L-J, Rafiq M, Radosa L, Hortschansky P, Cunha C, Cseresnyés Z, Krüger T, Schmidt F, Heinekamp T, Straßburger M, Löffler B, Doenst T, Lacerda JF, Campos A, Figge MT, Carvalho A, Kniemeyer O, Brakhage AA. 2023. Aspergillus fumigatus hijacks human p11 to redirect fungal-containing phagosomes to non-degradative pathway. Cell Host & Microbe 31:373–388.e10.

24. Uwamahoro N, Verma-Gaur J, Shen H-H, Qu Y, Lewis R, Lu J, Bambery K, Masters SL, Vince JE, Naderer T, Traven A. 2014. The Pathogen Candida albicans Hijacks Pyroptosis for Escape from Macrophages. mBio 5:e00003–14.

25. Yang L, Rojas AM, Shiau CE. 2021. Liposomal Clodronate-mediated Macrophage Depletion in the Zebrafish Model.

26. Schoen TJ, Rosowski EE, Knox BP, Bennin D, Keller NP, Huttenlocher A. 2019. Neutrophil phagocyte oxidase activity controls invasive fungal growth and inflammation in zebrafish. J Cell Sci 133:jcs236539.

27. Gratacap RL, Scherer AK, Seman BG, Wheeler RT. 2017. Control of Mucosal Candidiasis in the Zebrafish Swim Bladder Depends on Neutrophils That Block Filament Invasion and Drive Extracellular-Trap Production. Infect Immun 85:e00276–17.

28. Gratacap RL, Bergeron AC, Wheeler RT. 2014. Modeling Mucosal Candidiasis in Larval Zebrafish by Swimbladder Injection. J Vis Exp 52182.

29. Sharma S. 2016. Fungal Infection in Thermal Burns: A Prospective Study in a Tertiary Care Centre. JCDR 10.7860/JCDR/2016/20336.8445.

30. Calum H, Moser C, Jensen PØ, Christophersen L, Maling DS, van Gennip M, Bjarnsholt T, Hougen HP, Givskov M, Jacobsen GK, Høiby N. 2009. Thermal injury induces impaired function in polymorphonuclear neutrophil granulocytes and reduced control of burn wound infection. Clin Exp Immunol 156:102–110.

31. Deng Q, Yoo SK, Cavnar PJ, Green JM, Huttenlocher A. 2011. Dual roles for Rac2 in neutrophil motility and active retention in zebrafish hematopoietic tissue. Dev Cell 21:735–745.

32. Knox BP, Deng Q, Rood M, Eickhoff JC, Keller NP, Huttenlocher A. 2014. Distinct innate immune phagocyte responses to Aspergillus fumigatus conidia and hyphae in zebrafish larvae. Eukaryot Cell 13:1266–1277.

33. Ramakrishnan G, Miskolci V, Hunter M, Giese MA, Münch D, Hou Y, Eliceiri KW, Lasarev MR, White RM, Huttenlocher A. 2024. Real-time imaging reveals a role for macrophage protrusive motility in melanoma invasion. bioRxiv 10.1101/2024.09.25.614908.

34. Chen T, Wagner AS, Tams RN, Eyer JE, Kauffman SJ, Gann ER, Fernandez EJ, Reynolds TB. 2019. Lrg1 Regulates β (1,3)-Glucan Masking in Candida albicans through the Cek1 MAP Kinase Pathway. mBio 10:10.1128/mbio.01767-19.

35. Wagner AS, Lumsdaine SW, Mangrum MM, King AE, Hancock TJ, Sparer TE, Reynolds TB. 2022. Cek1 regulates ß(1,3)-glucan exposure through calcineurin effectors in Candida albicans. PLOS Genetics 18:e1010405.

36. Wagner AS, Hancock TJ, Lumsdaine SW, Kauffman SJ, Mangrum MM, Phillips EK, Sparer TE, Reynolds TB. 2021. Activation of Cph1 causes ß(1,3)-glucan unmasking in Candida albicans and attenuates virulence in mice in a neutrophil-dependent manner. PLOS Pathogens 17:e1009839.

37. Macherla C, Sanchez DA, Ahmadi MS, Vellozzi EM, Friedman AJ, Nosanchuk JD, Martinez LR. 2012. Nitric Oxide Releasing Nanoparticles for Treatment of Candida Albicans Burn Infections. Front Microbiol 3:193.

38. Sanchez DA, Schairer D, Tuckman-Vernon C, Chouake J, Kutner A, Makdisi J, Friedman JM, Nosanchuk JD, Friedman AJ. 2014. Amphotericin B releasing nanoparticle topical treatment of *Candida spp.* in the setting of a burn wound. Nanomedicine: Nanotechnology, Biology and Medicine 10:269–277.

39. Desai JV. 2018. Candida albicans Hyphae: From Growth Initiation to Invasion. 1. Journal of Fungi 4:10.

40. Davis SE, Hopke A, Minkin SC, Montedonico AE, Wheeler RT, Reynolds TB. 2014. Masking of β(1-3)- Glucan in the Cell Wall of Candida albicans from Detection by Innate Immune Cells Depends on Phosphatidylserine. Infect Immun 82:4405–4413.

41. Yang M, Solis NV, Marshall M, Garleb R, Zhou T, Wang D, Swidergall M, Pearlman E, Filler SG, Liu H. 2022. Control of β-glucan exposure by the endo-1,3-glucanase Eng1 in Candida albicans modulates virulence. PLOS Pathogens 18:e1010192.

42. Baltussen TJH, Zoll J, Verweij PE, Melchers WJG. 2019. Molecular Mechanisms of Conidial Germination in Aspergillus spp. Microbiology and Molecular Biology Reviews : MMBR 84:e00049.

43. de Jesus Carrion S, Leal SM, Ghannoum MA, Pearlman E. 2013. The RodA hydrophobin on Aspergillus fumigatus spores masks Dectin-1 and Dectin-2 dependent responses and enhances fungal survival in vivo. J Immunol 191:2581–2588.

44. Gravelat FN, Beauvais A, Liu H, Lee MJ, Snarr BD, Chen D, Xu W, Kravtsov I, Hoareau CMQ, Vanier G, Urb M, Campoli P, Abdallah QA, Lehoux M, Chabot JC, Ouimet M-C, Baptista SD, Fritz JH, Nierman WC, Latgé JP, Mitchell AP, Filler SG, Fontaine T, Sheppard DC. 2013. Aspergillus Galactosaminogalactan Mediates Adherence to Host Constituents and Conceals Hyphal β-Glucan from the Immune System. PLOS Pathogens 9:e1003575.

45. Scherer AK, Blair BA, Park J, Seman BG, Kelley JB, Wheeler RT. 2020. Redundant Trojan horse and endothelial-circulatory mechanisms for host-mediated spread of Candida albicans yeast. PLOS Pathogens 16:e1008414.

46. Tanner CD, Rosowski EE. 2024. Macrophages inhibit extracellular hyphal growth of A. fumigatus through Rac2 GTPase signaling. Infection and Immunity 92:e00380–23.

47. Culemann S, Knab K, Euler M, Wegner A, Garibagaoglu H, Ackermann J, Fischer K, Kienhöfer D, Crainiciuc G, Hahn J, Grüneboom A, Nimmerjahn F, Uderhardt S, Hidalgo A, Schett G, Hoffmann MH, Krönke G. 2023. Stunning of neutrophils accounts for the anti-inflammatory effects of clodronate liposomes. J Exp Med 220:e20220525.

48. Gur I, Zilbert A, Toledano K, Roimi M, Stern A. 2024. Clinical impact of fungal colonization of burn wounds in patients hospitalized in the intensive care unit: a retrospective cohort study. Trauma Surgery & Acute Care Open 9:e001325.

49. Horvath EE, Murray CK, Vaughan GM, Chung KK, Hospenthal DR, Wade CE, Holcomb JB, Wolf SE, Mason AD, Cancio LC. 2007. Fungal Wound Infection (Not Colonization) Is Independently Associated With Mortality in Burn Patients. Ann Surg 245:978–985.

50. Schofield CM, Murray CK, Horvath EE, Cancio LC, Kim SH, Wolf SE, Hospenthal DR. 2007. Correlation of culture with histopathology in fungal burn wound colonization and infection. Burns 33:341–346.

51. Pruitt BA, McManus AT, Kim SH, Goodwin CW. 1998. Burn wound infections: current status. World J Surg 22:135–145.

52. Gore AV, Monzo K, Cha YR, Pan W, Weinstein BM. 2012. Vascular Development in the Zebrafish. Cold Spring Harb Perspect Med 2:a006684.

53. Schoen TJ, Calise DG, Bok JW, Giese MA, Nwagwu CD, Zarnowski R, Andes D, Huttenlocher A, Keller NP. 2023. Aspergillus fumigatus transcription factor ZfpA regulates hyphal development and alters susceptibility to antifungals and neutrophil killing during infection. PLOS Pathogens 19:e1011152.

54. Gratacap RL, Rawls JF, Wheeler RT. 2013. Mucosal candidiasis elicits NF-κB activation, proinflammatory gene expression and localized neutrophilia in zebrafish. Disease Models & Mechanisms 6:1260.

55. Wu Y, Du S, Johnson JL, Tung H-Y, Landers CT, Liu Y, Seman BG, Wheeler RT, Costa-Mattioli M, Kheradmand F, Zheng H, Corry DB. 2019. Microglia and amyloid precursor protein coordinate control of transient Candida cerebritis with memory deficits. Nature Communications 10:58.

56. Schneider CA, Rasband WS, Eliceiri KW. 2012. NIH Image to ImageJ: 25 years of image analysis. Nat Methods 9:671–675.

57. Chen T, Jackson JW, Tams RN, Davis SE, Sparer TE, Reynolds TB. 2019. Exposure of Candida albicans β (1,3)-glucan is promoted by activation of the Cek1 pathway. PLOS Genetics 15:e1007892.

